# Rare ring conformations in PDB: Facts or wishful thinking?

**DOI:** 10.1101/2024.06.20.598822

**Authors:** Gabriela Bučeková, Viktoriia Doshchenko, Aliaksei Chareshneu, Jana Porubská, Michal Pajtinka, Michal Oleksik, Tomáš Svoboda, Vladimír Horský, Radka Svobodová

## Abstract

Protein structural data are highly valuable for research, and many significant results have been published on their basis. A key point for their credibility and applicability is their quality. An important facet of protein structure quality is the validation of ligands. Some aspects of ligand quality have already been validated by established quality metrics. However, validation of ring conformation has yet to be comprehensively performed despite rings strongly influencing the formation of the ligand’s scaffold and shape. Most rings form several conformations that differ in their stability. The most stable ones occur frequently in nature and should, therefore, be found in Protein Data Bank (PDB) structures.

In this article, we examined which conformations of rings occur in PDB structures. Our analysis focused on conformations of all cyclopentane, cyclohexane, and benzene rings in the PDB. Specifically, we examined 123 264 rings of 24 763 distinct ligands, which instances occur in 44 022 protein structures. In general, we found that most of the rings (98.32 %) are in energetically favourable conformations. Surprisingly, the existence of most of the energetically unfavourable ring conformations (2 067 samples, 1.68 %) is not supported by experimental data. Only 291 unfavourable ring conformations (0.24 %) are backed by experimental data that are accurate enough to distinguish the conformation, which shows that the existence of energetically unfavourable ring conformations is rarely supported by structural or experimental evidence.

Our results suggest that each occurrence of untypical ring conformation in the PDB may indicate a potential error and should be carefully analysed.

## Introduction

Experimentally determined protein structures, freely available in the Protein Data Bank (PDB) Berman et al. (2003), form a dataset of highly valuable research data. Many important results were published based on this data Lee et al. (2007); Waman et al. (2021); Batool et al. (2019); Knott et al. (2020). Moreover, these structures also serve as a training set for structure prediction of other currently experimentally unreachable proteins Jumper et al. (2021).

A key point for the credibility and applicability of this data is its quality, which is variable wwPDB consortium et al. (2019). Quality problems led to the retraction of articles focused on protein structure even from highly impacted journals Chang et al. (2006); Matthews (2007). For this reason, structure quality criteria have been developed Read et al. (2011); Henderson et al. (2012); Montelione et al. (2013); Trewhella et al. (2013), and validation reports have become a firm part of PDB Gore et al. (2017).

Many aspects of protein quality are validated and described by established quality metrics. For example, proper protein backbone torsion angles and torsion angles of amino acid side chains are evaluated by comparison to a high-quality reference dataset. Another validation approach searches for pairs of atoms that are impossibly close to each other in a structure model Gore et al. (2017). The agreement of the structure with its underlying experimental data is also validated Wang et al. (2022); Gore et al. (2017) despite the refinement process being somewhat subjective Kleywegt (2009).

Ligands, present in structure models alongside biomacromolecules, are essential for research in drug design and other pharmaceutical fields Scapin (2015). Just like their larger companions, these small molecules suffer from variable quality Deller and Rupp (2015), which stirred the scientific community to begin validating them Adams et al. (2016). That being said, the ligand validation field is still developing since not all aspects of ligands with fluctuating quality are being validated Nicklaus et al. (1995); Perola and Charifson (2004); Smart et al. (2018). The key components of ligands are rings, which strongly influence the formation of the ligand’s scaffold and shape. Most rings can form several conformations, which differ by their potential energy and, consequently, their stability. The most stable ones occur frequently in nature and, therefore, in ligands frequently found in structures in the PDB.

An important question is whether rings can occur only in stable conformations in protein structures or if they can form unfavourable conformations under certain conditions (e.g. when forced by the shape of the binding site). In our article, we focus on this unanswered question. Specifically, we selected a few common rings (cyclohexane, cyclopentane, and benzene) and analysed all their instances in the PDB. We determined their conformation and evaluated the occurrence of individual conformations.

We focus solely on ligand structures obtained using Xray crystallography since it is still the dominant structure determination method for complexes of biopolymers and ligands. More than 90 % of entries in PDB that contain at least one ligand, as well as more than 80 % of entries with an author-designated ligand of interest, were refined with this method as of 13th March 2024 (https://www.rcsb.org/stats/summary). The electron density available for the vast majority of entries enables us to assess whether the modelled conformation of a ligand is supported by its source experimental data.

## Methods

### Analysed rings

Our analysis focuses on three types of rings: cyclopentane, cyclohexane, and benzene. We selected these rings because they are the simplest representatives of the five- and six-membered rings. Known conformations of these rings are depicted in Figure 1.

**Figure 1.**
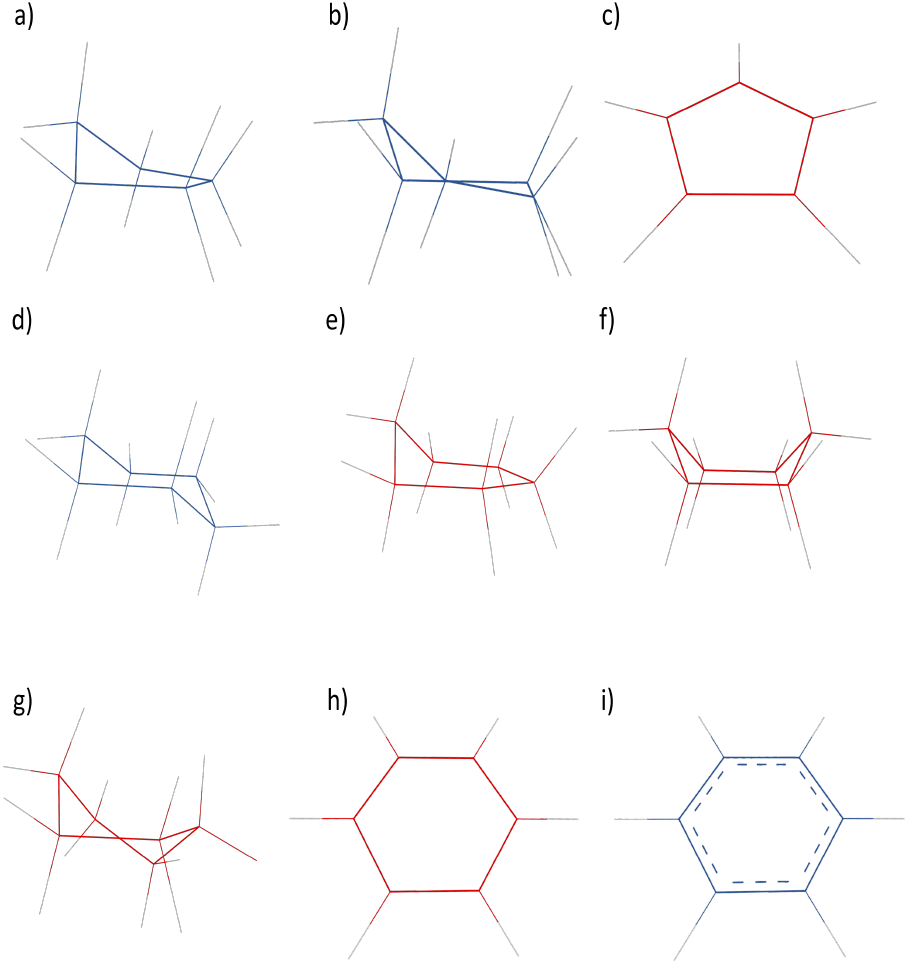
Conformation of analysed rings: Cyclopentane: a) envelope, b) half-chair, c) flat. Cyclohexane: d) chair, e) half-chair, f) boat, g) twist-boat, h) flat. Benzene: i) flat. Rings depicted in blue represent energetically favourable conformations, and rings illustrated in red represent rare conformations.

The conformation of molecules is responsible for their total energy. Bonds are limited in the extent of rotation in a ring, leading to different strains in a molecule. Cyclopentane can take on one of two low-energy conformations Tice et al. (2017): the envelope and the half-chair (as shown in Figure 1a and 1b). The envelope has four carbons in the plane, and the fifth atom is lifted. The second puckered conformation, the half-chair, is obtained by slight twisting of the envelope conformer, causing only three carbon atoms to remain coplanar, while carbons 3 and 5 are at different heights over the plane.

In the case of cyclohexanes, there are four known conformations Nelson and Brammer (2011): chair, half-chair, boat and twist-boat (as shown in Figure 1d to 1g). Out of these, the chair conformation is the most stable, with carbon 1 and 4 heading out of the plane in opposite directions. The boat conformation is less stable and is obtained by moving carbons 1 and 4 out of the plane in the same direction. Slightly more stable than the boat is twist-boat conformation obtained from the boat conformation by twisting two parallel carbon-carbon bonds against each other. The half-chair conformation has the highest energy among the four. The literature presents two versions of the half-chair conformation: one with five coplanar carbon atoms and one with four coplanar carbons. We consider only the former variant in this article.

Benzene naturally occurs only in the flat conformation (Figure 1i) due to the presence of delocalised electrons that form two p-orbitals below and above the ring and, along with *σ* electrons, enforce the benzene to adopt the regular hexagon shape with bond angles of 120° Vollhardt and Schore (2014).

Cyclopentane and cyclohexane rings have one more conformation: flat, where all atoms lie on a plane (Figure 1c and 1h). This conformation should only be hypothetical due to its high energy, therefore these rings should not acquire it.

### Workflow of the analyses

For the analysis of ring conformations, we used a workflow depicted in Figure 2. The analysis was performed on data acquired on 17th April 2024. During our analysis, we produced three datasets each one for every target ring type (i.e. cyclopentane, cyclohexane, and benzene datasets). The workflow consists of the following seven steps. All the software used in this workflow can be found in a GitHub repository (https://github.com/sb-ncbr/rings-conformation-validation), while the input data used for this analysis, the results, and a snapshot of the workflow are available at https://doi.org/10.58074/hy79-qc22.

**Figure 2.**
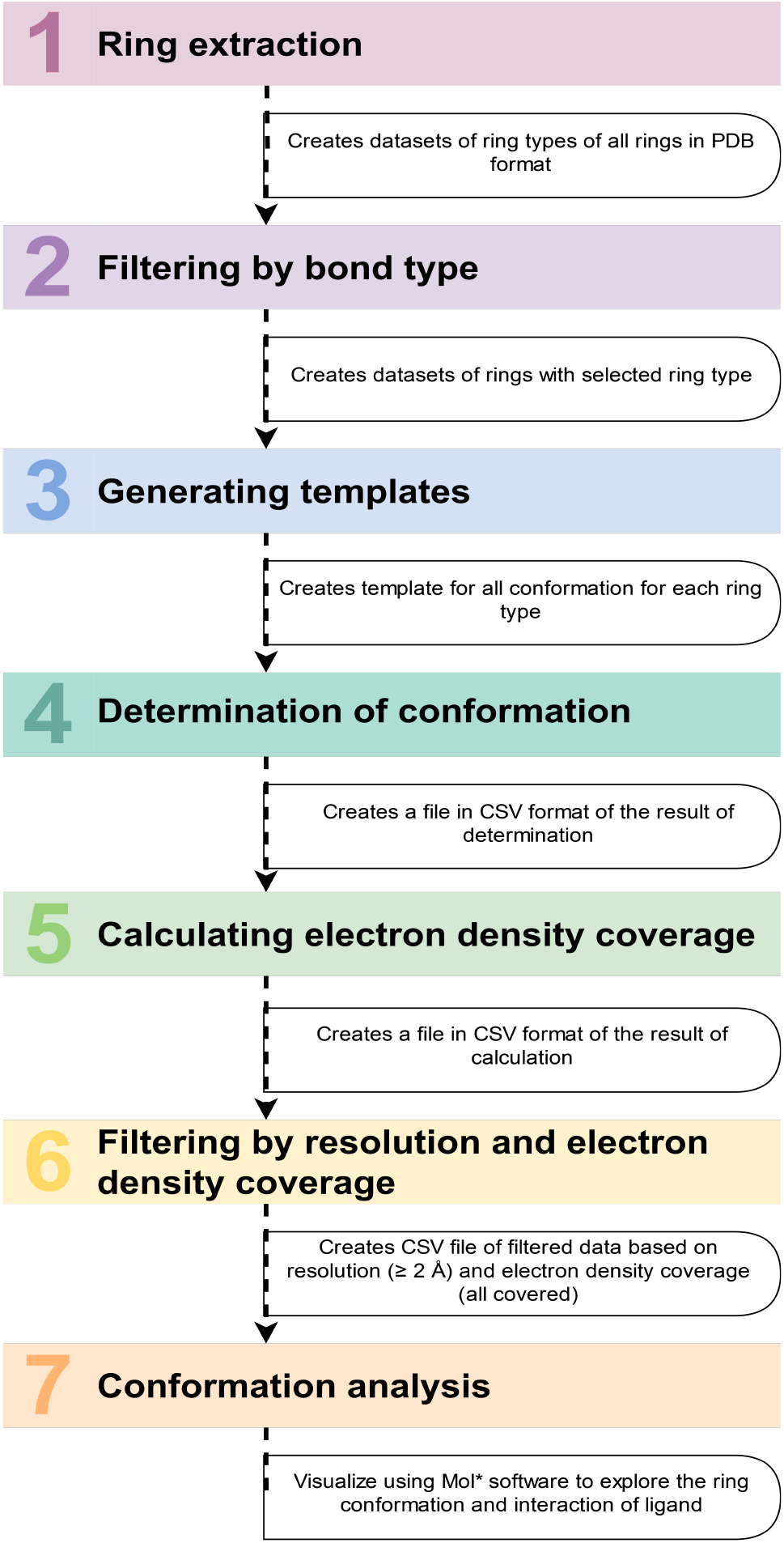
Schematic representation of conformational analysis workflow.

#### Ring extraction

As the first step, we used the Chemical Component Dictionary (CCD) Berman et al. (2003) in mmCIF format and selected ligands with at least one target ring. Subsequently, we extracted the rings from all instances of these ligands across the whole PDB via the PatternQuery tool Sehnal et al. (2015). As a result, we obtained each extracted ring as a separate PDB file, which we then sorted into datasets by the target ring types.

#### Filtering by bond type

PatternQuery uses its own query language, which, unfortunately, cannot distinguish between different bond types. Hence, our datasets required an additional filtration step to remove rings with unwanted types of bonds. For this step, we again used the CCD to get the information about bond types by mapping the atom names of our extracted rings in PDB files to corresponding atom names in the CCD. Based on that information, we discarded rings that were not cyclopentanes, cyclohexanes, or benzenes.

#### Generating templates

We determined the conformation of analysed rings by comparing their structure with templates of their conformations. The algorithm for generating such templates involved two steps. First, we divided PDB structures’ rings obtained in the previous step on the basis of their structure to produce as many subsets as there are different conformations for an analysed ring type. The model of each conformation was defined in the in-house program we used in this step of the workflow. The definition is based on identifying the central plane of the ring, which goes through the maximum number of atoms possible in a ring and divides this ring into up to three groups of atoms: those lying on the plane, over the plane, and under the plane.

Then, we superimposed all rings in each subset to compute an “average” ring. The superposition was computed by SiteBinder Sehnal et al. (2012). After discarding 10 % of the rings (rounded up) that were conformationally the most outlying on the basis of their root mean square deviation (RMSD) being the highest, we repeated the superimposition followed by discarding of outliers until only one ring remained, which we considered to be the template for its conformation.

#### Determination of conformation

We again used SiteBinder to superimpose each target ring structure and each template of the same ring type and to compute the RMSD of such superimposition. The most probable conformation of each target ring was determined straightforwardly: the superimposition of its template with the target ring produced the lowest RMSD.

Two different sets of templates were used to analyse benzene rings. Initially, the flat conformation benzene template was used, but it was observed that some rings had been assigned a flat conformation with an RMSD value of *≥* 0.1 Å. Therefore, the dataset was analysed using templates of cyclohexane conformations.

#### Calculating electron density coverage

The next step of the analysis focused on determining whether all atoms of target rings are fully covered by electron density. Data for this step, i.e. electron density maps, were obtained from the RCSB Berman (2000).

We computed the electron density in two steps. First, the algorithm determined the electron density value for the coordinate of the centre of an individual atom via trilinear interpolation of the eight nearest nodes on the electron density map. Then, the obtained interpolated value of electron density was compared with the threshold of the electron density isosurface. In our case, this threshold was set to 1.5 *σ* Lamb et al. (2015) as this value is one of the molecular viewers’ most commonly used values Sehnal et al. (2017, 2021). If the obtained electron density value was higher than the threshold, then the electron density covered the given atom. Otherwise, it was considered not covered. This process was repeated for each atom in a ring. Finally, the total number of covered atoms of a ring was yielded, which enabled us to filter out rings with at least one atom not covered by electron density.

The approach was implemented as a Python script. We used the Python library Gemmi Wojdyr (2022) to work with electron density data and to compute the trilinear interpolation.

#### Filtering by resolution and electron density coverage

We combined the results from the previous steps into a single CSV file. Then, we enriched the results with the electron density coverage and resolution value of the structure of each target ring. The resolution values were acquired from the PDBe using ValTrendsDB Horský et al. (2019) via data download. The obtained information was used to further filter target rings based on the resolution of their structure. We considered only structures with a resolution of 2 Å or better in the analysis since electron density maps with lower resolution tend to lack the information content needed for confident determination of ring conformation Wlodawer and Dauter (2017); Minor (2007). Then, the value of electron density coverage was assigned to individual analysed rings. To correctly determine the location and possible conformation, all atoms of the rings should be covered with electron density. Only fully covered rings were further analysed.

#### Conformation analysis

We selected a subset of rings with an untypical conformation from the results of the previous steps for subsequent data analysis. We used Mol* visualisation software to individually visualise ligands with rings from the subset and explore their position towards the protein structure and its possible interaction sites. A variety of case studies of untypical ring conformations are described further in the article.

### Limitations

During the analysis, we encountered a few limitations. Firstly, the determination of the conformation of the ring is sometimes equivocal due to the parameters necessary for the determination of the conformation being too benevolent or, on the contrary, strict. Therefore, conformations may overlap when the parameters are too benevolent, which may lead to the script assigning the first tested conformation to a target ring.

Secondly, identifying and extracting benzene that shares an edge with another benzene in polycyclic molecules like naphthalene and anthracene can be problematic. The single and double bonds alternate in this ring, which can result in the second benzene ring being incorrectly determined as cyclohexadiene.

## Results

We created three datasets while working on the analysis. The first dataset comprises 4 888 cyclopentane rings extracted from 856 different ligands. The second one with cyclohexanes contains 7 157 analysed rings from 1 653 ligand molecules. The last dataset includes 111 219 benzene rings extracted from 22 254 ligands. In total, we analysed 123 264 rings from 24 763 different ligand molecules. Detailed information about the datasets is summarised in Table 1.

**Table 1.** Representation of the target rings in the PDB. The three datasets consist of a total of 134 390 structures of the three target rings extracted from ligands.

|  | Number of distinct ligands in the CCD | Number of instances of ligands in the PDB | Number of analysed rings |
| --- | --- | --- | --- |
| Cyclopentane | 856 | 2 169 | 4 888 |
| Cyclohexane | 1 653 | 3 355 | 7 157 |
| Benzene | 22 254 | 38 498 | 111 219 |
| Total | 24 763 | 44 022 | 123 264 |

We used the workflow described in Section 2.2 to analyse the datasets. The results of the analyses are summed up in Table 2 and 3 and depicted in Figure 3. Additionally, templates of each conformation for the analysed ring types (except for a benzene) and CSV files containing the RMSD values, assigned conformation, resolution, and electron density coverage of each analysed ring are available at https://doi.org/10.58074/hy79-qc22.

**Table 2.** The overall statistics of the cyclopentane, cyclohexane, and benzene conformational analysis. The occurrence of individual conformations is shown for all analysed rings, for rings of ligands with a resolution of at least 2 Å, and for rings of ligands with a resolution of at least 2 Å with all ring atoms covered by electron density.

| Type of ring | Conformation | All rings | | Rings $\geq 2$ Å | | Rings $\geq 2$ Å and fully covered | |
| --- | --- | --- | --- | --- | --- | --- | --- |
|  |  | Number of ring | Occurs. [%] | Number of ring | Occurs. [%] | Number of ring | Occurs. [%] |
| Cyclopentane | Half-chair | 2 516 | 51.47 | 1 068 | 21.85 | 730 | 14.93 |
|  | Envelope | 1 952 | 39.93 | 775 | 15.86 | 467 | 9.55 |
|  | Flat | 420 | 8.59 | 172 | 3.52 | 71 | 1.45 |
|  | Total | 4 888 | 100.00 | 2 015 | 41.22 | 1 268 | 25.94 |
| Cyclohexane | Chair | 5 554 | 77.60 | 2 307 | 32.23 | 1 428 | 19.95 |
|  | Half-chair | 544 | 7.60 | 232 | 3.24 | 112 | 1.56 |
|  | Boat | 461 | 6.44 | 155 | 2.17 | 27 | 0.38 |
|  | Twist-boat | 534 | 7.46 | 213 | 2.98 | 54 | 0.75 |
|  | Flat | 64 | 0.89 | 30 | 0.42 | 15 | 0.21 |
|  | Total | 7 157 | 100.00 | 2 937 | 41.04 | 1 636 | 22.86 |
| Benzene | Flat | 111 175 | 99.96 | 50 594 | 45.49 | 31 525 | 28.35 |
|  | Chair | 17 | 0.02 | 14 | 0.01 | 10 | 0.01 |
|  | Half-chair | 24 | 0.02 | 3 | 0.00 | 2 | 0.00 |
|  | Boat | 2 | 0.00 | 0 | 0.00 | 0 | 0.00 |
|  | Twist-boat | 1 | 0.00 | 0 | 0.00 | 0 | 0.00 |
|  | Total | 111 219 | 100.00 | 50 611 | 45.51 | 31 537 | 28.36 |

**Table 3.** The statistical representation of the occurrence of energetically favourable and unfavourable conformations of the cyclopentane, cyclohexane, and benzene rings. The distribution of energetic conformation types is shown for all analysed rings, for rings of ligands with a resolution of at least 2 Å, and for rings of ligands with a resolution of at least 2 Å with all ring atoms being by electron density

| Type of ring | Conformation | All rings | | Rings $\geq 2$ Å | | Rings $\geq 2$ Å and fully covered | |
| --- | --- | --- | --- | --- | --- | --- | --- |
|  |  | Number of ring | Occurs. [%] | Number of ring | Occurs. [%] | Number of ring | Occurs. [%] |
| Cyclopentane | Energ. favr. | 4 468 | 91.41 | 1 843 | 37.70 | 1 197 | 24.49 |
|  | Energ. unfavr. | 420 | 8.59 | 172 | 3.52 | 71 | 1.45 |
|  | Total | 4 888 | 100.00 | 2 015 | 41.22 | 1 268 | 25.94 |
| Cyclohexane | Energ. favr. | 5 554 | 77.60 | 2 307 | 32.23 | 1 428 | 19.95 |
|  | Energ. unfavr. | 1 603 | 22.40 | 630 | 8.80 | 208 | 2.91 |
|  | Total | 7 157 | 100.00 | 2 937 | 41.04 | 1 636 | 22.86 |
| Benzene | Energ. favr. | 111 175 | 99.96 | 50 594 | 45.49 | 31 525 | 28.34 |
|  | Energ. unfavr. | 44 | 0.04 | 17 | 0.02 | 12 | 0.01 |
|  | Total | 111 219 | 100.00 | 50 611 | 45.51 | 31 537 | 28.36 |
| Summary | Energ. favr. | 121 197 | 98.32 | 54 744 | 44.41 | 34 150 | 27.70 |
|  | Energ. unfavr. | 2 067 | 1.68 | 819 | 0.66 | 291 | 0.24 |
|  | Total | 123 264 | 100.00 | 55 563 | 45.08 | 34 441 | 27.94 |

**Figure 3.**
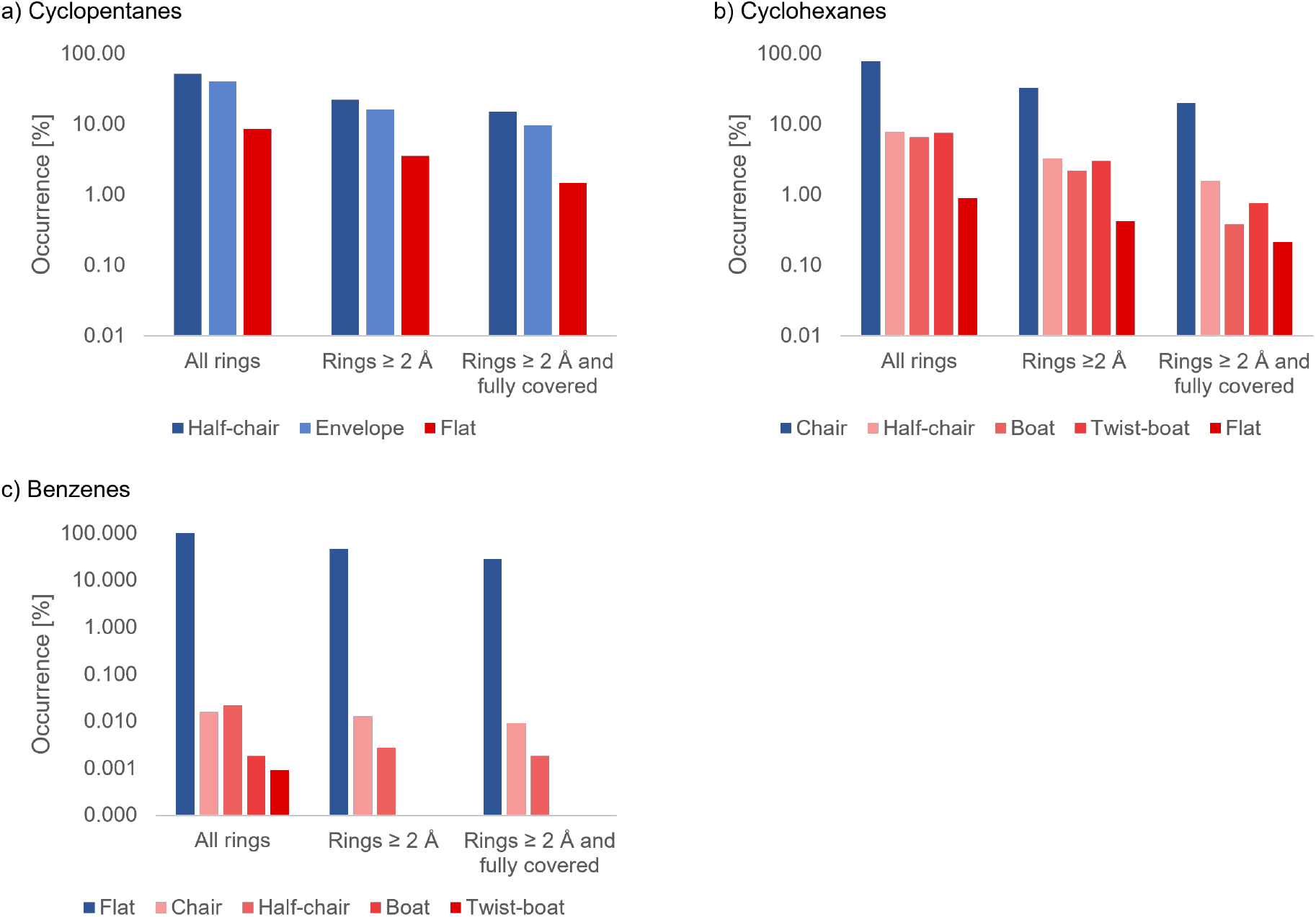
Occurrence of ring conformation of a) cyclopentanes, b) cyclohexanes, and c) benzenes. All histograms are on a logarithmic scale. Blue colour represents stable conformation for each type of ring. Red colour represents energetically unfavourable conformations.

### Analysis of the cyclopentane dataset

Upon thorough analysis of the cyclopentane dataset (Table 2 and 3), we have discovered that the CCD contains 856 unique ligands with at least one cyclopentane ring in their molecular structure. In the PDB, we found 4 888 instances of the rings in 2 169 individual protein structures. The cyclopentane rings in the dataset have three conformations, namely two stable conformations (half-chair and envelope) and one energetically unfavourable conformation (flat). Most of the rings were in stable conformations, and only about 8.59 % of cyclopentane rings (i.e. 420 rings) from the dataset were in flat conformation.

We further analysed the resolution and coverage of the target rings and found that only 71 of these flat rings have sufficient experimental data to detect their conformation, i.e. their resolution equals to or is higher than 2 Å, and all carbons in the cyclopentane ring are covered by electron density. Therefore, the remaining 349 rings seem to have been assigned the flat conformation without adequate proof of the existence of such conformation. We analysed these 71 rings manually to discover if there was some structural reason for this conformation. In almost all cases, the envelope or the half-chair conformations seem to fit the density data better. An example of a cyclopentane in the flat conformation, which should have a more energetically favourable conformation instead, is the ring in ligand with CCD ID 3KT in PDB entry 4r91 Caldwell et al. (2014).

### Analysis of the cyclohexane dataset

Our comprehensive evaluation of the cyclohexane dataset (see Table 2 and 3) revealed that 1 653 unique ligands in the CCD contain at least one cyclohexane ring. 7 157 instances of these rings can be found in 3 355 macromolecular structures in the PDB. Cyclohexane rings in the dataset occur in 5 conformations. The majority (more than 77.60 %) occurred in the chair conformation, which is the only stable one. The boat conformation was observed in 6.44 % of the dataset, while less than 1 % of the rings were in the flat conformation. The remaining cyclohexane rings were in the half-chair and twist-boat conformation, which are transition states Hendrickson (1961); Squillacote et al. (1975). However, only a small percentage of these rings (22.86 %) were found to meet the necessary conditions for further examination (i.e. resolution equal to or higher than 2 Å, and all atoms covered by electron density). Almost 3 % of the analysed rings differed from the expected chair conformation. The half-chair conformation was the most commonly represented rare cyclohexane conformation, with an occurrence of 1.56 %.

Upon analysing cases of rings occurring in the flat conformation in detail, we discovered that most of these rings should be present in one of the more energetically stable conformations. An example of such a potential conformation issue is a ligand with CCD ID OM3 in PDB entry 4cfb. However, there was a case where the flat conformation was most likely justified: a ligand with CCD ID FHR in PDB entry 7dnc Dai et al. (2022). We paid attention to this specific ligand in our case studies.

### Analysis of the benzene dataset

The dataset for benzene (as shown in Table 2 and 3) includes more than 22 254 unique ligands from the CCD. These ligands contain 111 219 instances of the benzene ring present in 38 498 macromolecular structures in the PDB, which is quite a lot even after filtering the dataset with the quality criteria (i.e. resolution equal to or higher than 2 Å, and all atoms covered by electron density). At first, we only examined the flat conformation of these rings since it is the only possible conformation according to the literature Vollhardt and Schore (2014). However, we observed higher RMSD values in the resulting tables, prompting us to analyse the conformations using the same set of templates we used for the conformational analysis of the cyclohexanes. Uncommon conformations are scarce in this dataset, with more than 28 % of rings meeting the criteria for further analysis (i.e. resolution equal to or higher than 2 Å, and all atoms covered by electron density) occurring in the flat conformation. Only ten benzenes occurred in the chair conformation and two in the half-chair conformation. Upon closer examination, we found that all rings for which this conformation was determined can occur in a typical flat conformation.

### Case studies of rings with untypical conformation

Several intriguing structures have been chosen for case studies to illustrate some of the energetically unfavourable conformations found in the PDB.

The structure of PDB entry 4v9c Wang et al. (2012) was determined using X-ray crystallography with a rather low resolution of 3.30 Å. The validation report of the entry shows several error values for qualitative metrics, such as clashscore. During the analysis, we identified a few areas where the model does not fit ideally to the electron density map. The structure contains eighteen neomycins, one (chain B, residue number 3163) incorporating a flat cyclohexane conformation (Figure 4a). The existence of this conformation is, in our opinion, unjustifiable since the ligand’s target ring is not adequately covered by electron density, and several positive (i.e. green) density blobs near it indicate where the atom should be located.

**Figure 4.**
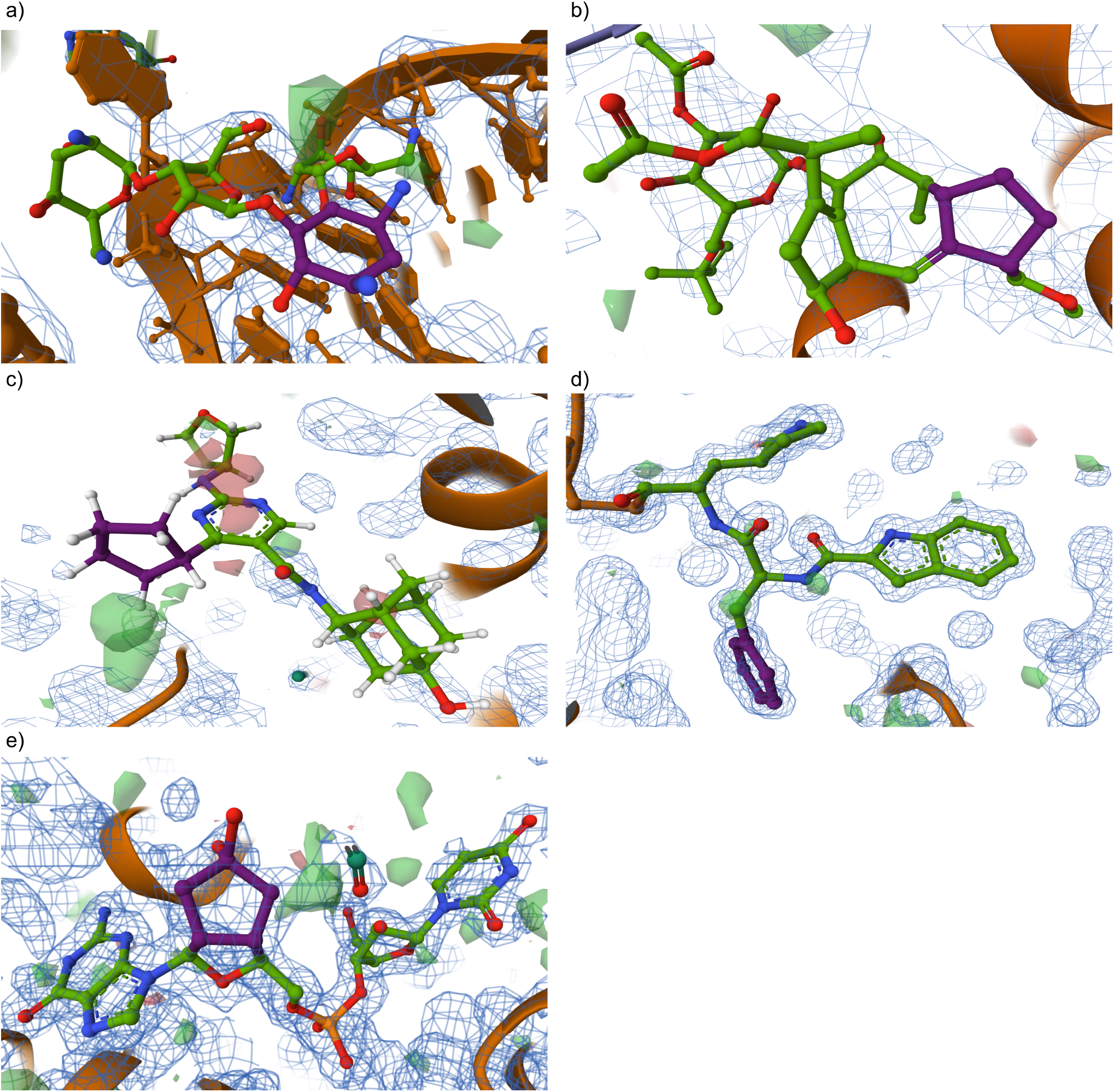
Visualisations of ligands from the PDB from the case studies. The Mol* visualisation software Sehnal et al. (2021) was used to visualise ligands with electron density maps. The blue mesh blobs represent the 2Fo-Fc *σ* map, and the green mesh blobs illustrate the Fo-Fc(+ve) *σ* and red mesh blobs illustrate the Fo-Fc(-ve) *σ* difference map. a) the neomycin (chain B, residue number 3163) in PDB entry 4v9c Wang et al. (2012) with analysed ring marked in purple, b) the fusicoccin (chain B, residue number 301) in PDB entry 5nwk Saponaro et al. (2017) with analysed ring marked in purple, c) the ligand HD2 (chain A, residue number 1551) in PDB entry 5alu Öster et al. (2015) with analysed ring marked in purple, d) the ligand FHR (chain A, residue number 201) in PDB entry 7dnc Dai et al. (2022) with analysed ring marked in purple, e) the ligand FGO (chain A, residue number 505) in PDB entry 7c19 Chandravanshi et al. (2021) with analysed ring marked in purple.

Subsequently, we analysed cyclopentane rings, which are present in fusicoccin molecules in PDB entry 5nwk Saponaro et al. (2017), which was released in 2017 and is determined at a resolution of 3.30 Å. Some qualitative factors of the structure that are present in its validation report have problematic values, such as clashscore or R-free Chen et al. (2010); Brünger (1992). The structure contains seven molecules of fusicoccin, differing in their electron density coverage. Of the seven cases where the cyclopentane ring is in the flat conformation (one is shown in Figure 4b), only four are entirely covered in electron density. It is important to note that fusicoccin is a flexible molecule that can change its conformation depending on its interactions with its surroundings Stevers et al. (2016); Skwarczynska et al. (2013). However, various studies of fusicoccin did not confirm the occurrence of flat conformation of cyclopentane in the ligand Ballio et al. (1991). Moreover, the low resolution of the structure makes it difficult to verify the correctness of the conformation of cyclopentane present in the structure.

PDB entry 5alu Öster et al. (2015) contains three instances of ligand with CCD identifier HD2, one of which (chain A, residue number 1 551) contains a cyclopentane in the flat conformation (Figure 4c). Remarkably, the ring lacks electron density coverage, while the other two ligands comprise an entirely covered cyclopentane ring in an energetically favourable conformation. Although the structure has a high resolution of 1.87 Å, it is difficult to support the presence of the flat conformation of the cyclopentane due to the lack of electron density coverage.

The next case study was conducted on planar cyclohexane, which can be found as part of the ligand with CCD identifier FHR (shown in Figure 4d) in PDB entry 7dnc Dai et al. (2022), released quite recently in 2021. The resolution of the structure is high at 1.17 Å. The validation report of the structure shows its good quality. The ligand, which has an inhibitor function, is located in the substrate-binding site in the macromolecular structure. All atoms of the ligand are covered by electron density. Although the flat conformation of the cyclohexane in question may not be energetically favourable when considering the ligand as an individual structure, there is a high probability of the occurrence of the flat conformation being connected to the biological function or interactions with residues of the macromolecule.

The last case study revolves around a cyclopentane in the structure of a ligand with CCD identifier FGO (chain A, residue number 505, shown in Figure 4e) in PDB entry 7c19 Chandravanshi et al. (2021). The target ring discussed is in the flat conformation. This structure 7c19 was released in 2021 with a high resolution of 1.77 Å. The cyclopentane in question is covered with electron density along with the entire ligand. The ring shares an edge with an oxolane on one side while having two hydroxy groups bound to an atom on the opposite side of the target ring. These hydroxy groups interact with the binding site of the macromolecule. The structural and interaction data support the planar conformation of the cyclopentane ring.

## Conclusion

We analysed conformations of all cyclopentane, cyclohexane, and benzene rings occurring in the PDB entries determined via X-ray crystallography. Specifically, we examined 123 264 rings originating from 24 763 distinct ligands, which instances occur in 44 022 protein structures.

Overall, we found that most rings (98.32 %) are in energetically favourable conformations. Surprisingly, the existence of most of the unfavourable ring conformations (2 064 samples, 1.68 %) we found is not supported by experimental data. Either the resolution of the structure is not high enough (i.e. is lower than 2 Å), or the ring atoms are not covered by electron density. We believe there is no reasoning in such cases to model rings to have untypical, energetically unfavourable ring conformations.

Only 291 unfavourable ring conformations (0.24 %) from the whole dataset are supported by experimental data. We analysed all such cases to determine if there is some structural reason for the untypical conformations to occur. The analysis showed us only a few cases where a ring adopting an untypical conformation makes sense.

The results of our work lead to the recommendation that each occurrence of an untypical ring conformation in a protein structure may indicate a site of concern or a potential error and should be carefully examined.

## Acknowledgements

Biological Data Management and Analysis Core Facility of CEITEC Masaryk University, funded by ELIXIR CZ research infrastructure (MEYS Grant No: LM2023055), is gratefully acknowledged for supporting the research presented in this paper. Computational resources were provided by the e-INFRA CZ project (ID:90254), supported by the Ministry of Education, Youth and Sports of the Czech Republic, and by the ELIXIR-CZ project (ID:90255), part of the international ELIXIR infrastructure. Persistent data storage was provided by the EGI DataHub (https://datahub.egi.eu) service, powered by Onedata (https://onedata.org) software - an open-source, distributed data management platform. We gratefully acknowledge Polish high-performance computing infrastructure PLGrid (HPC Centers: ACK Cyfronet AGH) for providing computer facilities and support within computational grant no. PLG/2023/016142 and the EGI-ACE project (grant number 101017567).

## Funding information

Czech Science Foundation [22-30571M]; Ministry of Education, Youth and Sports of the Czech Republic: ELIXIR CZ [LM2023055]. Funding for open access charge: Masaryk University as per its Read and Publish agreement with the Oxford University Press.

## Supplementary Information

All the software used in this workflow can be found in a GitHub repository (https://github.com/sb-ncbr/rings-conformation-validation). The input data used for this analysis, the results, and a snapshot of the workflow, as well as templates of each conformation for all three analysed ring types and CSV files containing the RMSD values, assigned conformation, resolution, and electron density coverage of each analysed ring are available at https://doi.org/10.58074/hy79-qc22.

## Notes

https://github.com/sb-ncbr/rings-conformation-validation

https://doi.org/10.58074/hy79-qc22

### Competing Interest Statement

The authors have declared no competing interest.

